# The *Bacillus subtilis* cell envelope stress-inducible *ytpAB* operon modulates membrane properties and contributes to bacitracin resistance

**DOI:** 10.1101/2024.01.17.576085

**Authors:** Jessica R. Willdigg, Yesha Patel, Briana E. Arquilevich, Chitra Subramanian, Matthew W. Frank, Charles O. Rock, John D. Helmann

**Author notes:** Address correspondence to: John D. Helmann. deceased (9.22.23) (Charles O. Rock).

## Abstract

Antibiotics that inhibit peptidoglycan synthesis trigger the activation of both specific and general protective responses. σ^M^ responds to diverse antibiotics that inhibit cell wall synthesis. Here, we demonstrate that cell wall inhibiting drugs, such as bacitracin and cefuroxime, induce the σ^M^-dependent *ytpAB* operon. YtpA is a predicted hydrolase previously proposed to generate the putative lysophospholipid antibiotic bacilysocin (lysophosphatidylglycerol), and YtpB is the branchpoint enzyme for the synthesis of membrane-localized C_35_ terpenoids. Using targeted lipidomics we reveal that YtpA is not required for the production of lysophosphatidylglycerol. Nevertheless, *ytpA* was critical for growth in a mutant strain defective for homeoviscous adaptation due to a lack of genes for the synthesis of branched chain fatty acids and the Des phospholipid desaturase. Consistently, overexpression of *ytpA* increased membrane fluidity as monitored by fluorescence anisotropy. The *ytpA* gene contributes to bacitracin resistance in mutants additionally lacking the *bceAB* or *bcrC* genes, which directly mediate bacitracin resistance. These epistatic interactions support a model in which σ^M^-dependent induction of the *ytpAB* operon helps cells tolerate bacitracin stress, either by facilitating the flipping of the undecaprenyl-phosphate carrier lipid or by impacting the assembly or function of membrane-associated complexes proteins involved in cell wall homeostasis.

**Importance:** Peptidoglycan synthesis inhibitors include some of our most important antibiotics. In *Bacillus subtilis*, peptidoglycan synthesis inhibitors induce the σ^M^ regulon, which is critical for intrinsic antibiotic resistance. The σ^M^-dependent *ytpAB* operon encodes a predicted hydrolase (YtpA) and the enzyme that initiates the synthesis of C_35_ terpenoids (YtpB). Our results suggest that YtpA is critical in cells defective in homeoviscous adaptation. Further, we find that YtpA functions cooperatively with the BceAB and BcrC proteins in conferring intrinsic resistance to bacitracin, a peptide antibiotic that binds tightly to the UPP lipid carrier that sustains peptidoglycan synthesis.

## Introduction

The bacterial cell envelope, minimally consisting of a plasma membrane and a peptidoglycan cell wall, is the major barrier between the cell interior and the extracellular environment (1). Peptidoglycan is a rigid and highly cross-linked structure that confers cell shape and resists turgor pressure to prevent lysis. It is also the target of some of the most clinically relevant antibiotics (2). When exposed to changing environmental conditions, including antibiotics, bacteria respond through dedicated cell envelope stress response (CESR) pathways and modulate gene expression to protect the integrity of the cell envelope (3, 4).

Following exposure to a cell envelope stress, an extracellular signal must be communicated across the cell envelope to mediate a transcriptional response. Bacterial CESRs are commonly controlled by two component system (TCS) regulatory networks or by alternative σ factors (often members of the extracytoplasmic function or ECF σ family) which are in turn regulated by stress-responsive anti-σ factors (4–6). For example, the *Bacillus subtilis* BceRS TCS responds selectively to the peptide antibiotic bacitracin that binds to undecaprenyl-pyrophosphate (UPP) and inhibits the recycling of lipid II (7, 8). Activation of the BceRS system upregulates BceAB, an ABC transporter and a primary determinant of intrinsic bacitracin resistance (8). BceAB acts through a target protection mechanism to release bacitracin from inhibited UPP:bacitracin complexes (9). The BceAB complex also collaborates with the BceS sensor kinase to respond to bacitracin stress in a flux-sensing mechanism (10, 11).

In addition to TCSs, ECF σ factors play a prominent role in the regulation of CESRs (4, 6, 12). Upon sensing a cell envelope stress, the membrane-embedded anti-σ factor is inactivated (often by proteolytic cleavage) to release the active σ factor (13–16). One such ECF σ, σ^M^, is activated by cell wall targeting antimicrobials (17–20), although the precise σ^M^ activating stimulus is unclear. The activation of σ^M^ induces the expression of nearly 100 genes, many of which are involved in peptidoglycan synthesis, cell shape determination, and cell division (6, 18). Consistently, deletion of *sigM* sensitizes the cell to β-lactams (21), moenomycin (20), bacitracin (17), and other antibiotics that target peptidoglycan synthesis (22).

The large σ^M^ regulon includes several operons with poorly characterized roles in responding to envelope stress (18). The *ytpAB* operon is one such example. Previously, the *ytpA* gene product was assigned as a class A_2_-phospholipase that cleaves the *sn2* acyl chain from phosphatidylglycerol (PG) to produce a discrete lysophospholipid (1-(12-methyltetradecanoyl)-3-phosphoglyceroglycerol). This lysophospholipid has been named bacilysocin, and was suggested to function as an antibiotic to inhibit growth of neighboring microorganisms (23). However, purified bacilysocin has weak antibiotic activity with an MIC for *Saccharomyces cerevisiae* of 5 µg/ml and 25 µg/ml for *Staphylococcus aureus* (23), and there is no evidence that it is released into the media at levels sufficient to demonstrate antibiotic activity.

The second gene in the operon, *ytpB*, encodes an enzyme required for sesquarterpenoid synthesis (tetraprenyl-β-curcumene synthase) (24, 25). Sesquarterpenoids are 35 carbon (C_35_) cyclic compounds derived from heptaprenyl-pyrophosphate (HPP) and have a multi-ring structure similar to C_30_ hopanoids, which are derived from squalene (26). The major sesquarterpenoid made by *B. subtilis* is designated baciterpenol A (24), which can be further modified by autooxidation and dehydration (under non-physiological isolation conditions) to generate baciterpenol B and sporulenes (27). Deletion of *ytpB* leads to a modest increase in cell sensitivity towards bacitracin (28). Our previous findings revealed that this effect results from increased accumulation of the YtpB substrate, HPP (28). The linear C_35_ HPP isoprenoid is a structural analog of the longer C_55_ isoprenoid, UPP, and both contain a membrane-proximal pyrophosphate moiety, which is the ligand for bacitracin (29).

Here, we have explored the role of YtpA, a putative phospholipase, on membrane properties and bacitracin sensitivity. Overexpression of *<u>ytpA</u>* increased membrane fluidity, but in contrast with a prior report (23), YtpA was not required for lysophosphatidylglycerol (LPG) production. Genetic studies reveal that *ytpA* is critical for the fitness of cells defective in homeoviscous adaptation. Moreover, YtpA contributes to bacitracin resistance in parallel with the BceAB and BcrC resistance systems. We propose that YtpA may support peptidoglycan synthesis by modulating membrane properties to enhance the function of the synthetic machinery and perhaps to facilitate the transmembrane flipping of the UP carrier lipids (30).

## Results

### Overexpression of YtpA increases membrane fluidity

In previous studies, YtpA was identified as a lysophospholipase responsible for synthesis of bacilysocin (1-(12-methyltetradecanoyl)-3-phosphoglycerol (1-15-LPG)), a lysophospholipid derived from phosphatidylglycerol with a 15 carbon anteiso branched chain fatty acid (23). In eukaryotes, lysophospholipids participate in membrane remodeling via the Lands’ cycle in which phospholipids can be deacylated and then reacylated with chemically distinct fatty acids (31). However, the presence of this type of membrane remodeling pathway in bacteria has yet to be established (32). If lysophospholipids persist within the membrane bilayer, they may alter local membrane curvature, permeability, and fluidity (33–35).

To determine if YtpA impacts membrane fluidity we used the membrane intercalating dye 1,6-diphenyl-1,3,5-hexatriene (DPH) to perform fluorescence anisotropy (FA) (36). The rotational freedom of DPH in the membrane serves as an indicator of membrane fluidity. There was no significant difference in FA between the *B. subtilis* 168 (*trpC2*) wild-type strain (WT) and an isogenic *ytpA* null strain (Δ*ytpA*) (Figure 1). However, since the *ytpAB* operon is known to be stress-inducible (18) this may simply mean that *ytpA* is poorly expressed under these growth conditions. Therefore, we tested the effect of YtpA overexpression using an IPTG-inducible promoter. Indeed, expression of *ytpA* led to a decrease in FA compared to the WT and Δ*ytpA* strain (Figure 1). This suggests an increase in rotational freedom, which is indicative of an increase in membrane fluidity (36). In a previous study, DPH measurements of FA in vesicles made from *B. subtilis* membrane lipids revealed a near linear decrease in FA (an increase in fluidity) over the temperature range from 10 °C to 45 °C (37). The change observed here between WT cells and those with induction of *ytpA* is comparable to vesicles incubated at temperatures differing by 15-20 °C (37). Further, a similar magnitude of change was seen in *B. subtilis* cells with and without induction of the σ^W^-dependent membrane stress response, which significantly protects cells against detergents and other agents that increase fluidity (38). Thus, the effect seen here is likely to be physiologically significant.

**Figure 1.**
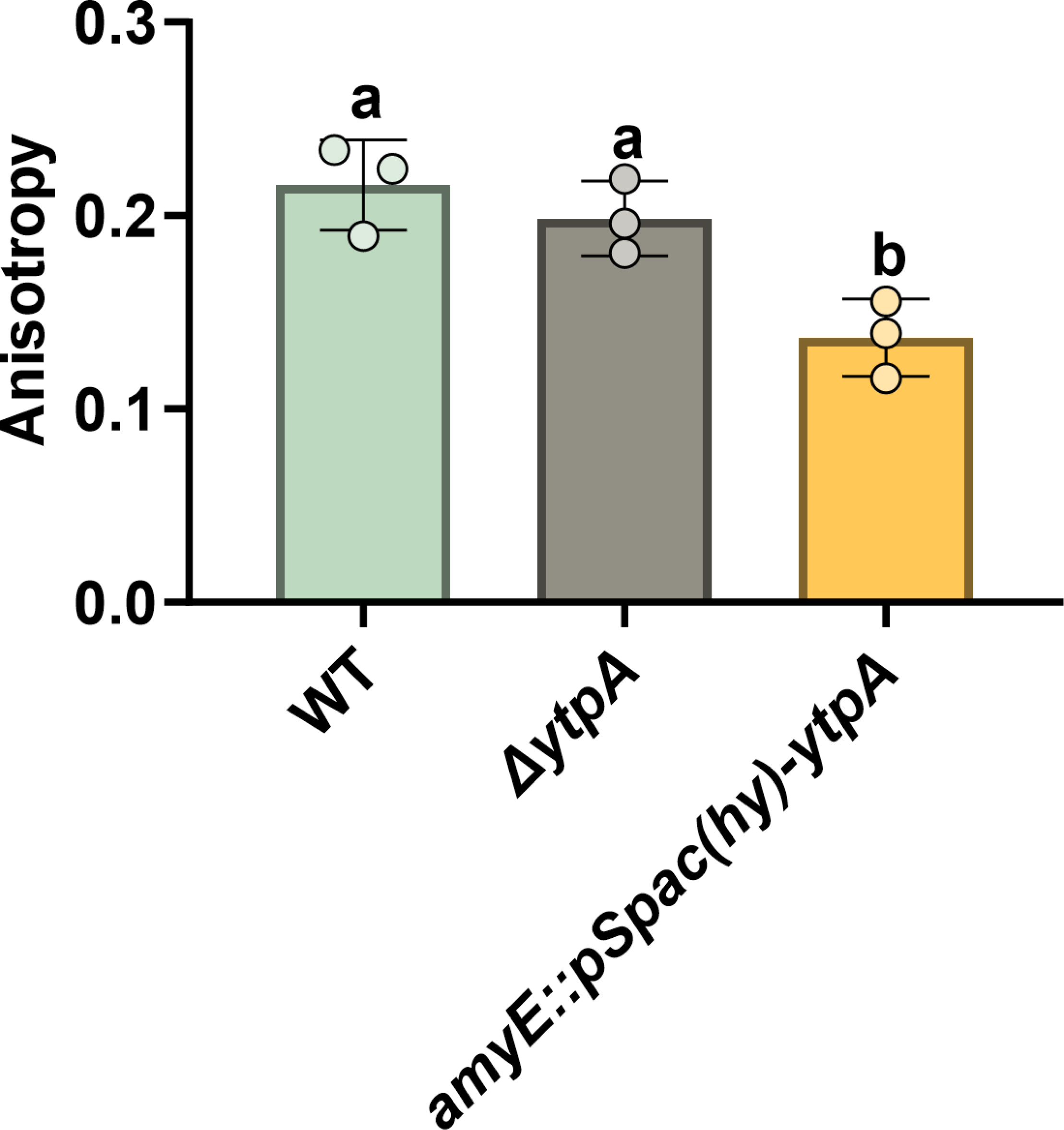
Induction of *ytpA* increases membrane fluidity. Overexpression of *ytpA* using the *spac(Hy)* promoter (HB27450) with 1 mM IPTG yields statistically significant differences in anisotropy compared to the WT and *ytpA* knockout (Δ*ytpA,* HB27232) strains. N = 3 biological replicates. A one-way ANOVA with a Tukey test for multiple comparisons was performed. Columns labeled with different letters are statistically distinct from each other; with a *P* value cutoff < 0.05.

### Induction of *ytpA* rescues growth of cells defective in homeoviscous adaptation

To test if the membrane fluidizing effect noted upon induction of *ytpA* is physiologically relevant, we took advantage of a reporter strain with an artificially rigid membrane (39). This strain, designated Δ*bkd*, is defective in homeoviscous adaptation to conditions of low fluidity due to deletions of the *bkd* operon and the *des* gene. These mutations prevent the synthesis of branched chain fatty acids (BCFA) and the desaturation of acyl chains by the Des desaturase, respectively (39). Approximately 90% of the *B. subtilis* membrane is composed of BCFAs that help confer an optimal fluidity necessary for the maintenance of the electron transport chain (40). As a result, the Δ*bkd* strain requires supplementation with precursors to BCFAs for normal growth. For example, supplementing with 2-methylbutyric acid (MB) restores the ability to synthesize *anteiso* BCFAs and rescues growth in minimal medium (39).

The Δ*bkd* strain has a minor growth defect compared to WT when grown on LB medium at 27 °C, 37 °C and 45 °C. However, on deletion of *ytpA* (Δ*bkd* Δ*ytpA*), fitness was dramatically reduced. The colony size of the Δ*bkd* Δ*ytpA* strain was very small compared to both the WT and Δ*bkd* strain under all the temperatures tested (Figure 2A). As previously reported, the Δ*bkd* strain is inviable at 22 °C on minimal medium lacking BCFA precursors (Figure 2B). Remarkably, Δ*bkd* with a copy of *ytpA* expressed from the *spac(Hy)* promoter is able to grow at 22 °C, and if MB is additionally present this strain grows as well as WT (Figure 2B). The rescue of growth by YtpA is observed both with and without addition of the inducer IPTG, consistent with the known leaky expression of the *spac(Hy)* promoter (41). Using real time PCR we estimate that the leaky expression from this promoter leads to a two-fold increase in gene expression compared to its native expression in the WT cells. These results suggest that *ytpA* expression is critical to compensate for the growth-limiting defects that define the Δ*bkd* strain.

**Figure 2.**
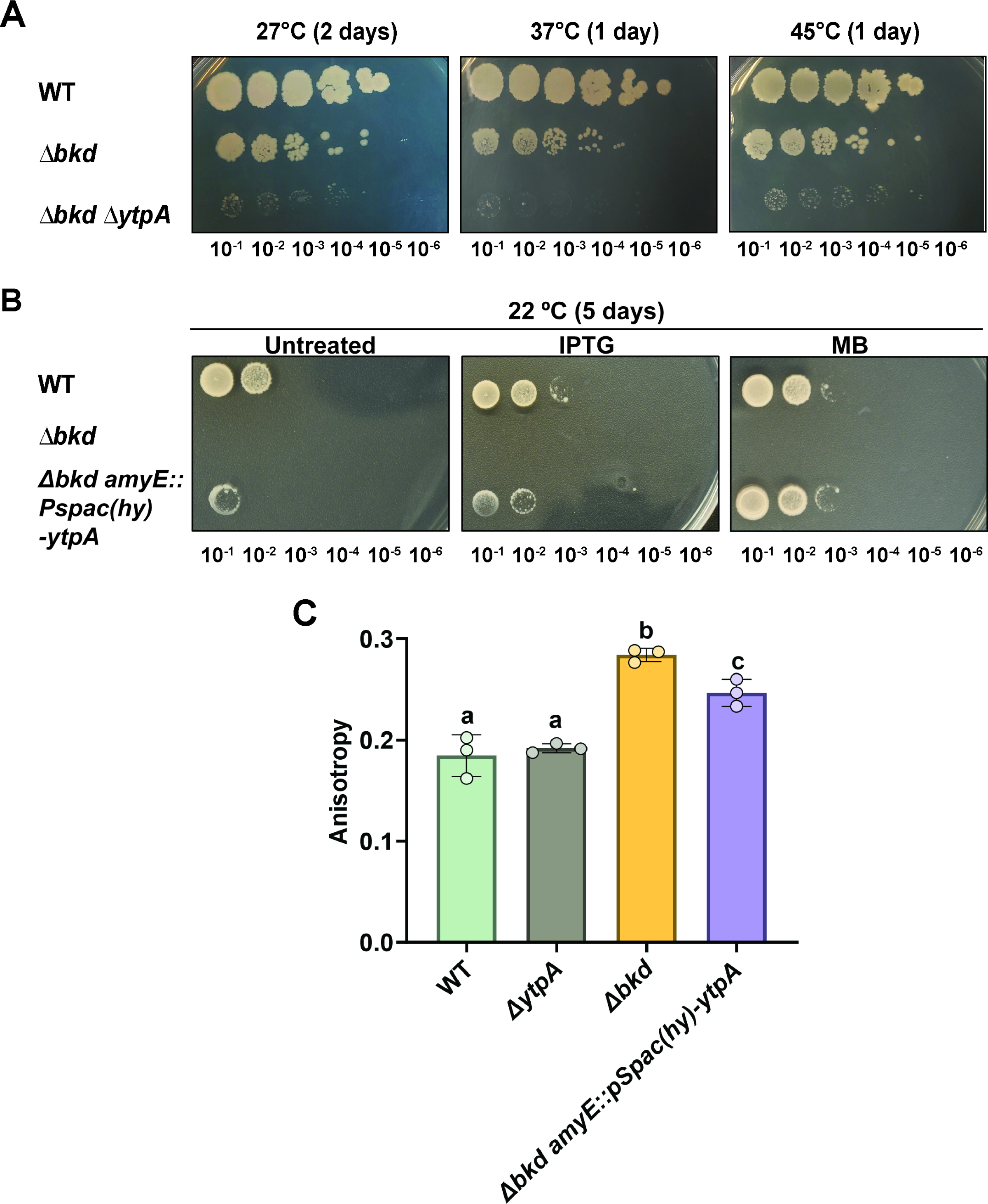
YtpA is physiologically important in cells with defects in membrane fluidity (A). A clean, unmarked deletion of *ytpA (*Δ*ytpA)* in the Δ*bkd* strain (HB27482), reduces the growth of the cells at the permissive temperatures of 27°C, 37 °C and 45 °C on LB medium. The colony size of the Δ*bkd* Δ*ytpA* strain is significantly smaller than the Δ*bkd* strain (HB27373). (B) Overexpression of *ytpA* in Δ*bkd* cells (HB27384) restores viability on minimal media when grown at non-permissive low temperature (22 °C). A representative image is shown (N = 3). Untreated column represents cells plated on minimal media without any supplementation. IPTG column represents cells plated on minimal media supplemented with 1 mM IPTG and MB column represents cells plated on minimal media supplemented with 100 μM 2-methylbutyric acid. (C) Induction of *ytpA* from the IPTG-inducible *spac(Hy)* promoter partially restores fluidity in a *Δbkd* strain. The data presented is the average of three biological replicates where errors bars represent the standard deviation. A one-way ANOVA with a Tukey test for multiple comparisons was performed. Columns labeled with different letters are statistically distinct from each other with a *P* value cutoff of < 0.05.

We next used FA to test if induction of YtpA increases membrane fluidity of the Δ*bkd* strain (Figure 2C). As reported previously (39), the Δ*bkd* strain shows an increase in FA compared to WT. Induction of *ytpA* in the Δ*bkd* strain led to a significant decrease in FA, although not a complete restoration back to the levels of WT cells (Figure 2C). This is consistent with the partial rescue of growth by induction of *ytpA* in the Δ*bkd* strain at 22 °C in the absence of MB (Figure 2B). We conclude that expression of YtpA increases membrane fluidity and restores growth of a strain defective in biochemical pathways that normally serve to increase membrane fluidity.

### YtpA is not the major phospholipase *in vivo*

YtpA was proposed to be a phospholipase A_2_ responsible for the release of 1-(12-methyltetradecanoyl)-3-phosphoglycerol (1-15-LPG) into the medium (23). Because lysophospholipids may impact membrane biology, we performed a targeted lipidomic analysis to determine if deletion of *ytpA* altered the phospholipid and lysophospholipid content of the cell. In both the WT and *ytpA* mutant strain (Δ*ytpA*), the major phosphatidylglycerol (PG) and phosphatidylethanolamine (PE) species were the same, with the most abundant species having a total of 30, 31, or 32 carbons in the acyl chains, with a C_15_ fatty acid in the 2-position. The minor 28-PG/PE and the 29-PG/PE peaks have a C_13_ or C_14_ acyl chain in the 2-position, respectively (Supplementary Figure 1A and 1B).

Next, we analyzed the lysophosphatidyglycerol (LPG) and lysophosphatidylethanolamine (LPE) composition of cells (Figure 3 and Supplementary Figure 1C, 1D, 2). For quantitation, the signals arising from the 1- and 2-acyl-lysophospholipids with the same carbon number were combined. Note that bacilysocin, the previously described C_15_ 1-acyl-lysophospholipid (23), most likely results from the action of an unidentified A_1_ phospholipase (producing a 2-acyl-lysophospholipid as the product) followed by fatty acyl chain migration (Supplementary Figure 1), which reaches ∼90% at the 1-position at equilibrium (42–44). Similar acyl chain migration was seen in recent studies monitoring lysophospholipid production in *Staphylococcus aureus* (45).

**Figure 3.**
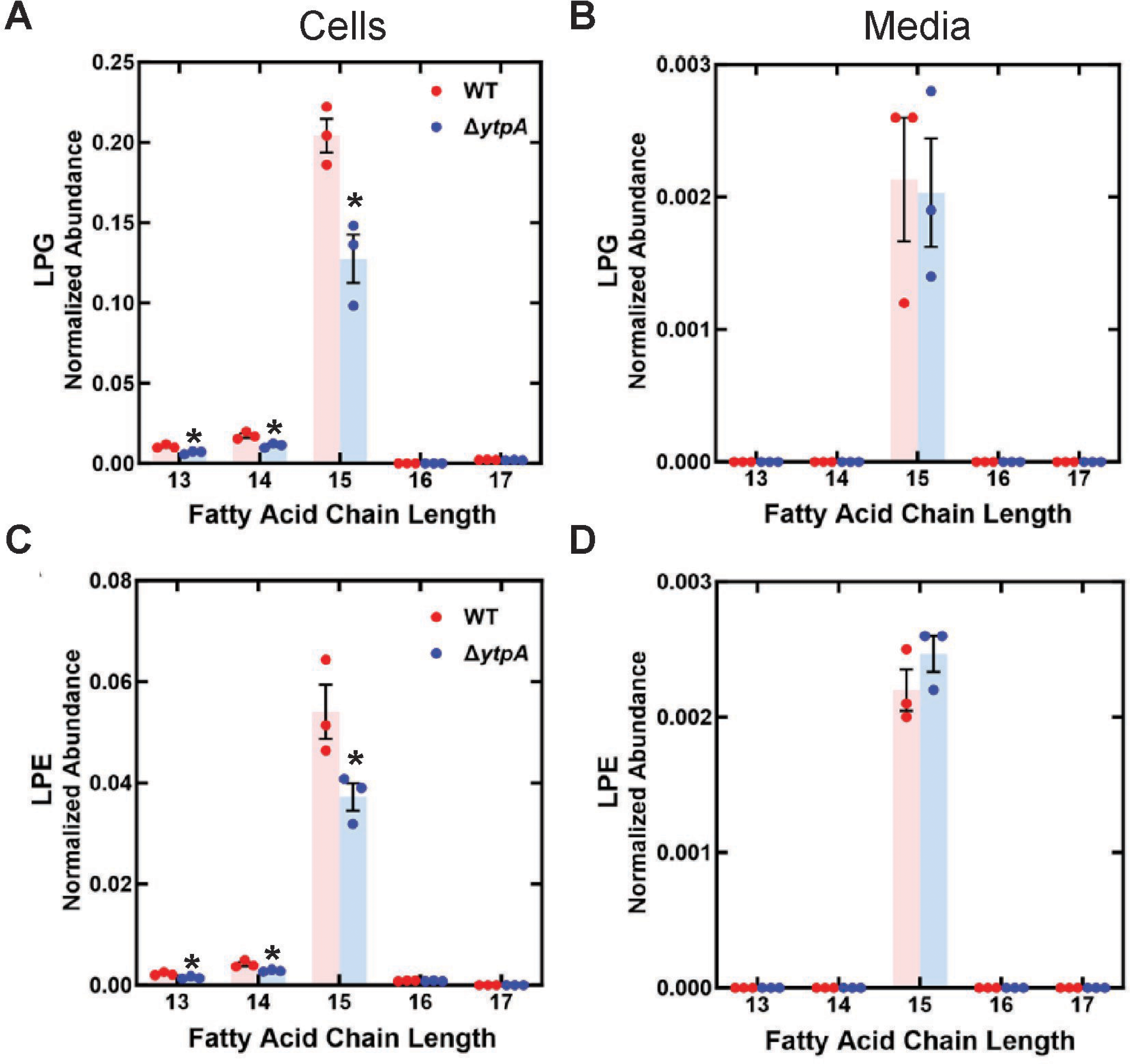
Lysophospholipid content of cells and media in WT and Δ*ytpA* strain (HB27232). Strains were grown in LB to late-log phase, the cells or media were extracted with methanol and the LPG/LPE molecular species determined by LC-MS/MS. The LPG/LPE abundances were determined relative to a [d5]17-LPG internal standard. WT (red); Δ*ytpA* (blue). (A) cellular LPG, (B) media LPG, (C) cellular LPE, (D) media LPE. Student t-test was done to compare the values of WT and *ytpA* samples for each molecular species separately. * indicates *P* value < 0.05.

In growing cells and in the medium, the dominant lysophospholipid was 15-LPG (Figure 3A, 3B), with minor amounts of 13- and 14-LPG in cells (Figure 3A). 15-LPE was also a major species in both cells (Figure 3C) and media (Figure 3D) in late-log phase. There was a modest, but statistically significant, reduction in lysophospholipids in the Δ*ytpA* strain in growing cells (Figure 3A, 3C). There was little if any effect noted in stationary phase cells (Supplementary Figure 2A, 2C). One notable change in the stationary phase cultures was that the amount of lysophospholipids in the medium was significantly elevated (10-fold) compared to samples taken in late-log phase (Supplementary Figure 2B, 2D). The presence of lysophospholipids in the cellular fraction of the Δ*ytpA* strain suggests that YtpA is not responsible for the bulk of lysophospholipid synthesis in *B. subtilis.* Consistently, induction of *ytpA* with IPTG did not result in an increase in lysophospholipids compared to either the uninduced condition or WT (Supplementary Figure 3).

YtpA is a member of a large superfamily of serine-dependent hydrolases (alpha-beta hydrolases) with a wide variety of substrates. Bioinformatic searches indicate that YtpA is likely a cytosolic-facing, membrane-associated protein. Sequence homology searches consistently yield sequence and domain similarities between YtpA and other phospholipases, including PldB, a poorly characterized phospholipase and the namesake of the COG2267 superfamily. Because traditional homology searches are limited to sequence similarity, we additionally used the YtpA AlphaFold2-generated structure to search protein structure databases using FoldSeek (46–48). Amongst the many proteins with similar predicted structures, biochemical information is available for only a handful. For example, YtpA has 30% identity to a secreted monoacylglycerol hydrolase from *Mycobacterium tuberculosis* (UNIPROT 007427) that hydrolyses glycerol monoesters of long-chain fatty acids (49, 50). The lack of a clear functional role for YtpA highlights a recurring problem for this large family of alpha/beta hydrolases, enzymes that often still have enigmatic functions (51).

### Cell envelope active antibiotics induce expression of *ytpAB* in a σ^M^-dependent manner

Next, we evaluated the regulation of *ytpA* in conditions that lead to cell envelope stress. The *ytpAB* operon is regulated by σ^M^, an alternative ECF sigma factor that is activated in response to various peptidoglycan synthesis inhibitors and other cell wall stressors (6). Early genome-wide transcriptome studies have revealed that *ytpAB* is most strongly induced by inhibitors of the membrane-associated steps of peptidoglycan biosynthesis, and in particular by those compounds that interfere with the lipid II cycle such as bacitracin and vancomycin (18, 52). This pattern of response is consistent with a recent comprehensive profiling study using both RNA-seq and tiling array methodologies in which bacitracin and vancomycin were the strongest inducers, followed by tunicamycin, moenomycin, and lysozyme (53).

We constructed a luciferase transcriptional reporter to monitor the expression of the *ytpAB* operon in response to cell envelope stresses. Consistent with expectation, the *ytpAB* reporter fusion was strongly induced by high levels of bacitracin (31.25 μg/ml), and this induction was lost if either the σ^M^ promoter site or the *sigM* gene was deleted (Figure 4). The reporter fusion was also induced by cefuroxime (0.16 μg/ml), a drug that inhibits the activity of enzymes involved in peptidoglycan synthesis. Using real-time PCR, we observed a four-fold induction of *ytpA* after 15 min of treatment with 31.25 μg/ml of bacitracin, and a two-fold increase when cells were grown to mid-log phase in the presence of the same concentration of bacitracin.

**Figure 4.**
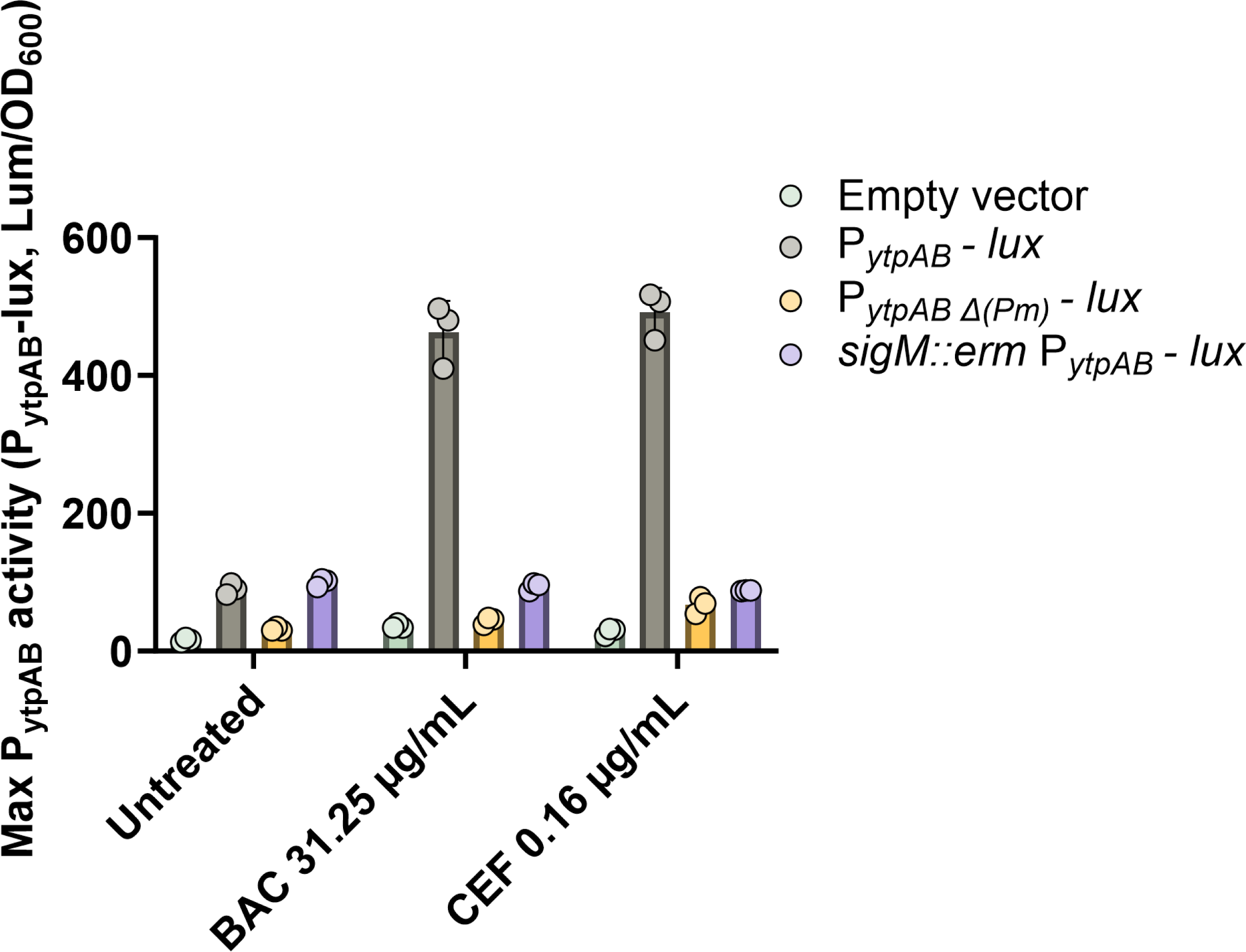
Induction of the *ytpAB* operon is σ^M^ dependent. Induction of *PytpAB-lux* (HB27246) is lost following bacitracin treatment in the absence of *sigM* (HB27287) and the σ^M^-specific consensus sequence at the *ytpAB* promoter (HB27247). N = 3 biological replicates; error bars represent standard deviation.

Deletion of bacitracin resistance genes (*bcrC, bceAB*) significantly sensitizes the cell to bacitracin and is known to alter the expression of bacitracin-responsive genes (7). While a WT cell had a bacitracin MIC of 125 µg/ml, deletion of the intrinsic bacitracin resistance determinants *bceAB* or *bcrC* significantly reduced the MIC. In a *bceAB* or *bcrC* deletion background, *ytpAB* expression was induced by concentrations of bacitracin as low as 1.25 µg/ml, (Supplementary Figure 4). The observation that *ytpA* expression is induced in response to cell wall acting drugs is suggestive of a role in intrinsic drug resistance.

### The *ytpAB* operon confers bacitracin resistance

Next, we sought to determine if the *ytpAB* operon contributes to bacitracin resistance. The individual deletions of *ytpA* (Δ*ytpA*) or *ytpB* (Δ*ytpB*) did not have a significant effect on the growth of the cells with 62.5 µg/ml bacitracin (0.5x MIC). However, the *ytpAB* double mutant (Δ*ytpAB*) had a notable growth lag (Figure 5). An effect of YtpB on bacitracin resistance was noted previously in studies in Mueller-Hinton medium, and was ascribed to the accumulation of the YtpB substrate heptaprenyl-pyrophosphate (HPP), a close chemical analog of UPP (28). HPP likely complexes with bacitracin and may reduce the efficiency of BceAB-dependent detoxification by competition for the active site of the BceAB resistance protein (9). HPP might also serve as a competitive substrate for the BcrC-dependent phosphatase (28). Although Δ*ytpB* did not affect bacitracin resistance under the conditions we tested (LB medium), *ytpA* or *ytpB* together clearly contribute to bacitracin resistance.

**Figure 5.**
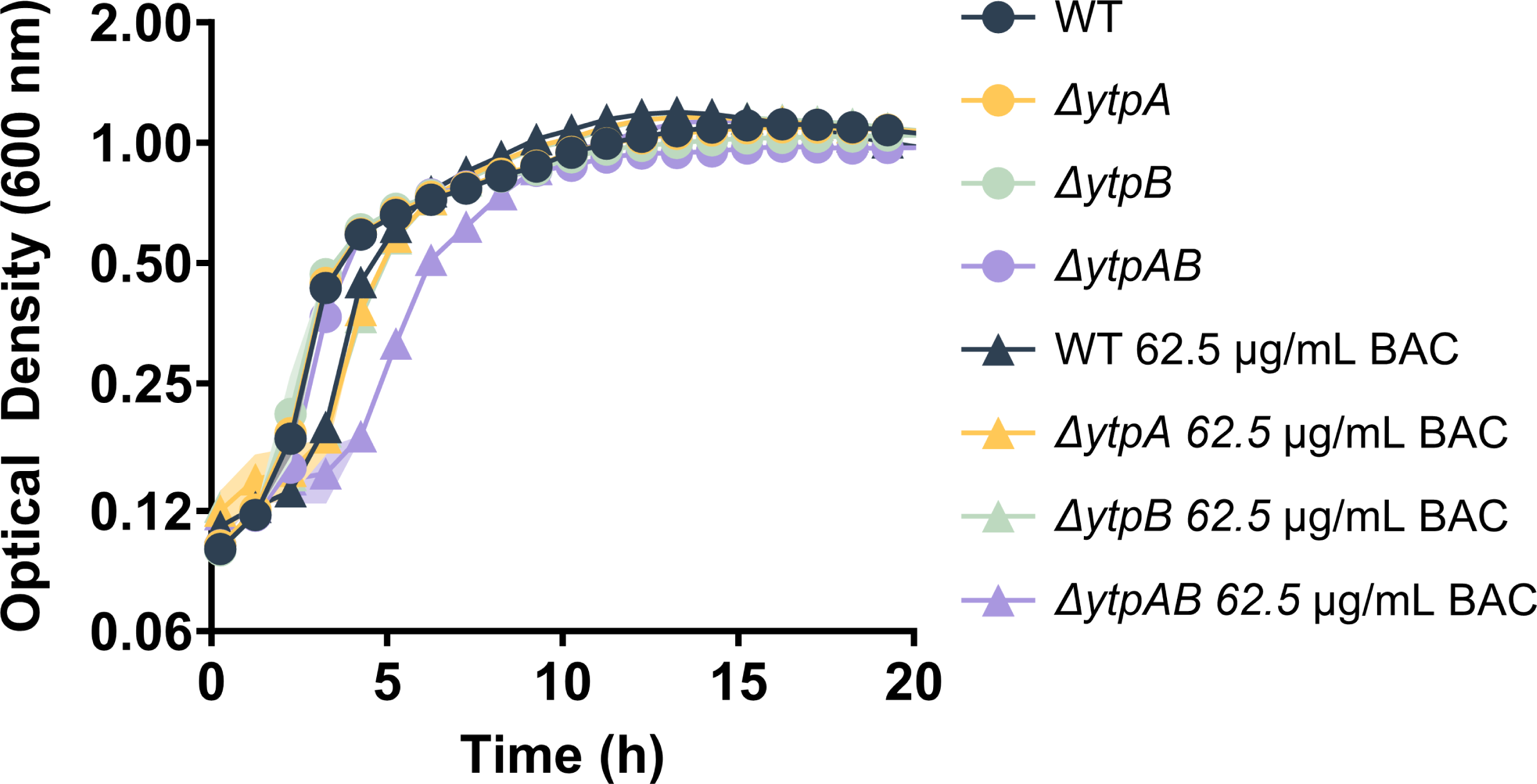
Effects of Δ*ytpA* and Δ*ytpB* (unmarked, in-frame deletions) on bacitracin sensitivity. Individual deletions of *ytpA* (HB27232) or *ytpB* (HB27253) do not have a significant effect on bacitracin (BAC) sensitivity compared to the WT cells. The *ytpAB* operon deletion (HB27407) has an increased lag in the presence of 62.5 μg/mL bacitracin compared to either single mutant. N =6; standard deviation in the growth of each strain has been shown by shading.

### Loss of YtpA increases bacitracin sensitivity in strains lacking BcrC or BceAB

We reasoned that if YtpA were contributing to bacitracin resistance by increasing membrane fluidity, one mechanism might be through facilitation of UP (or UPP) flipping across the membrane. To explore this, we monitored the bacitracin sensitivity of strains defective in recycling of the undecaprenyl carrier lipid due to lack of a UPP phosphatase (BcrC or UppP). The BcrC and UppP phosphatases are individually dispensable, but the double mutant is not viable (54, 55). In the presence of low levels of bacitracin (5 μg/ml) neither the Δ*bcrC* nor the Δ*ytpA* mutants displayed much of a growth lag. However, the Δ*ytpA* Δ*bcrC* double mutant was greatly inhibited with a >4 hr growth lag (Figure 6A). In contrast, there is very little additivity between Δ*ytpA* and Δ*uppP*, even with high bacitracin levels (62.5 μg/ml) (Figure 6B).

**Figure 6.**
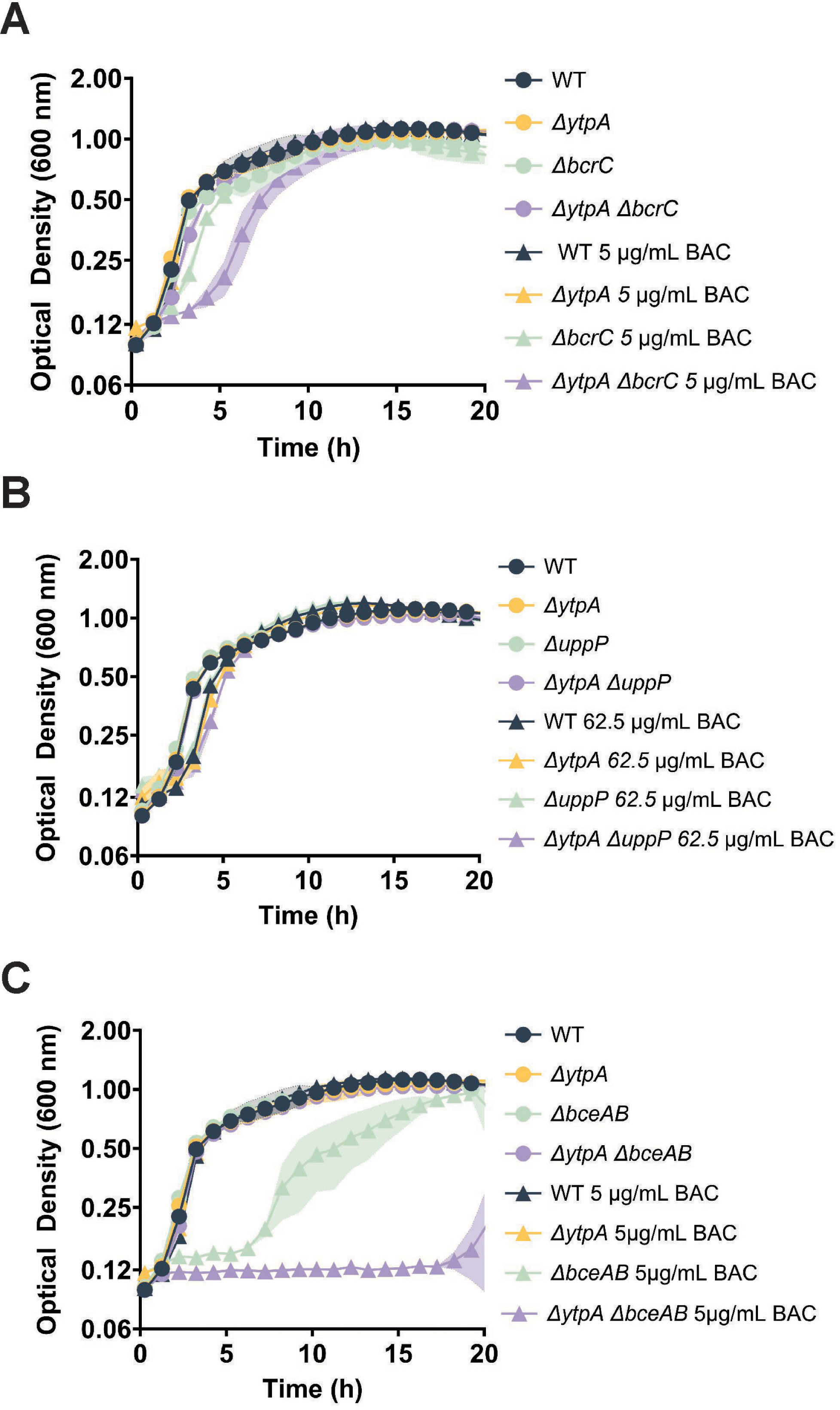
Loss of YtpA increases bacitracin (BAC) sensitivity in genetically sensitized strains. (A) Deletion of *ytpA* (HB27360) increases the sensitivity of the bacitracin-sensitive *bcrC* mutant (HB27277) as measured with 5 μg/ml bacitracin (B) Neither the loss of *ytpA* (HB27232) or *uppP* (HB27446) alone, nor the combination (HB27447), has a major impact on bacitracin sensitivity as measured with 62.5 μg/ml bacitracin. (C) Deletion of *ytpA* (HB27273) increases the sensitivity of the bacitracin-sensitive *bceAB* mutant (HB27271) as measured with 5 μg/ml bacitracin. N = 5 biological replicates; standard deviation in the growth of each strain has been shown by shading.

Next, we explored the role of YtpA in strains lacking the BceAB resistance pathway. The BceAB proteins function in the dissociation of UPP:bacitracin complexes in a target protection mechanism of resistance (9). Consistent with prior work (9), Δ*bceAB* was highly sensitive to bacitracin with decreased growth observed at 5 μg/ml. At this concentration, the Δ*ytpA* strain was unaffected, wherease the Δ*ytpA* Δ*bceAB* double mutant was unable to grow (Figure 6C). The additivity of YtpA with both BcrC and BceAB, the two major players of bacitracin resistance network, suggest a role for YtpA in bacitracin resistance. By increasing membrane fluidity, YtpA may reduce UPP levels on the outer leaflet of the membrane, perhaps by allowing it to flip inside.

### Loss of YtpA has only a small effect in strains lacking the UptA UP flippase

Following the transglycosylation reaction, the UPP lipid carrier is dephosphorylated on the outer leaflet of the membrane. Then, the transmembrane flipping of the UP product is facilitated by DedA family membrane proteins (22, 56). In *B. subtilis*, the σ^M^-regulated *uptA* (formerly *yngC* gene) encodes one such protein (22). A null mutant of *uptA* (Δ*uptA*) has no overt growth defect but displays an increased sensitivity to MX-2401 (22), an antibiotic that binds selectively to UP exposed on the outer leaflet of the membrane (57).

We speculated that YtpA-dependent membrane changes might also help to support the flipping of UP. However, deletion of *ytpA* did not increase the sensitivity of the *uptA* strain (Δ*ytpA* Δ*uptA*) to the sub-MIC level of 0.6 μg/ml MX-2401 (Figure 7A). Consistent with the notion that UptA mediates flipping of UP but not UPP, the *uptA* deletion (Δ*uptA*) had little effect on bacitracin sensitivity (Figure 7B), as shown previously (22). Moreover, the *ytpA* and *uptA* mutations did not exhibit an additive effect on bacitracin sensitivity (Figure 7B). The absence of additivity with UptA on sensitivity to MX-2401 suggests that YtpA has no significant role in modulating UP levels on the outside of the membrane.

**Figure 7.**
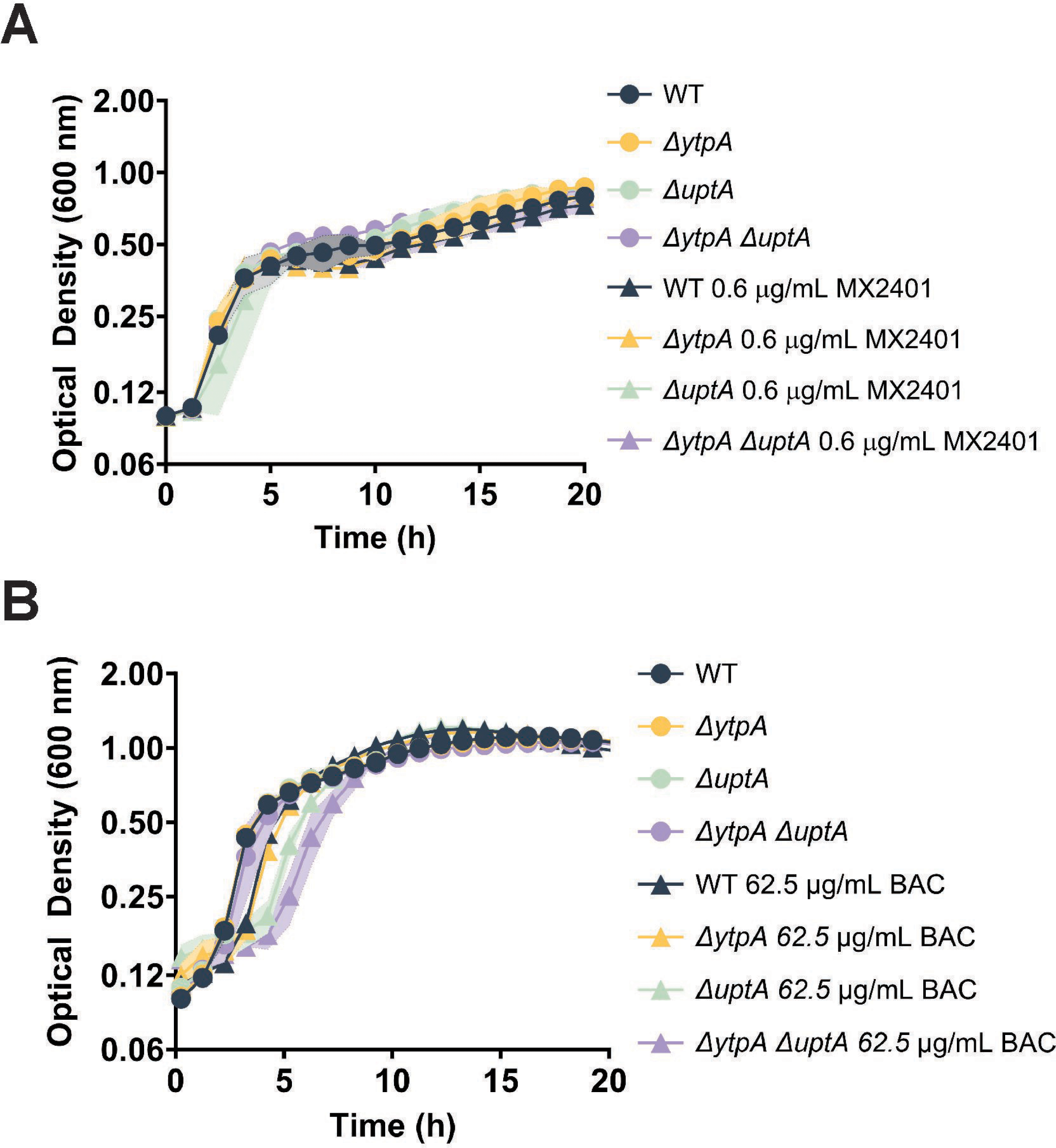
Epistasis of *ytpA* and *uptA* as measured by the growth of the *ytpA* (HB27232), *uptA* (HB27393) mutants alone and in combination (HB27362) on treatment with (A) 0.6 μg/ml MX-2401 and (B) 62.5 μg/ml bacitracin. N = 5 biological replicates; standard deviation in the growth of each strain has been shown by shading.

## Discussion

Antibiotics that interfere with peptidoglycan synthesis activate a large regulon of genes associated with σ^M^-dependent promoters that collectively function to sustain cell wall synthesis even if one or more steps are inhibited (6, 18). The σ^M^ stress response is triggered when the membrane-localized anti-σ^M^ complex (YhdK/YhdL) is inactivated by a still unknown mechanism (13). Induction is amplified by a very strong positive autoregulation that leads to high level but transient expression from an autoregulatory promoter for the *sigM-yhdL-yhdK* operon (58). Prolonged and un-regulated induction of the σ^M^ regulon is lethal due, in part, to toxicity from high level production of numerous integral membrane proteins (59). The transient induction of the σ^M^ stress response can counteract the action of many cell wall-acting antibiotics, and *sigM* mutants display heightened sensitivity to moenomycin, bacitracin, β -lactams, and other peptidoglycan synthesis inhibitors (20, 21).

To define the roles of σ^M^-activated operons in protection against cell envelope stress we and others have sought to identify σ^M^ target genes and their functions. This task is complex due to the large number of σ^M^-activated operons (18) and the overlapping regulation with other ECF σ-dependent regulons (19–21, 60). In addition, many stress-induced operons, including those for essential genes, are expressed independent of σ^M^ and then further upregulated in times of stress. The role of σ^M^ in protecting against peptidoglycan synthesis inhibitors can be attributed, at least in part, to the upregulation of genes for peptidoglycan synthesis. The σ^M^ regulon includes genes for both cytosolic steps of peptidoglycan biosynthesis (Ddl, MurB, MurF) and membrane-associated steps, including the alternate lipid II flippase (Amj), components of the Rod complex for peptidoglycan assembly (RodA, MreBCD), and a class A PBP (PBP1) (18, 61–63). In the specific case of moenomycin, σ^M^ regulation of the RodA transglycosylase is sufficient for resistance (61, 63). Other σ^M^-activated functions include enzymes for UPP synthesis (IspD, IspF) (64), the BcrC UPP-phosphatase (18, 60, 65), and the UptA UPP flippase (22), which can all function to help sustain sufficient levels of the undecaprenyl-phosphate lipid carrier (7, 64). In addition, σ^M^ increases synthesis of stress-induced isozymes for synthesis of lipoteichoic acid (LtaSa; (66)) and wall teichoic acid (TagT;(67)). Finally, σ^M^ activates genes that control secondary stress responses. The latter include genes encoding the SasA(YwaC) small alarmone synthase responsible for generation of (p)ppGpp, pGpp, ppApp, and AppppA (68), the DisA synthase for cyclic-di-AMP, and Spx, a transcription factor that controls a large regulon that contributes to protection from oxidative stress (69).

Although the roles of many σ^M^-regulated operons have now been defined, there are still others with poorly understood functions. Here, we demonstrate that the *ytpAB* operon contributes to the high level of intrinsic bacitracin resistance characteristic of *B. subtilis*. Bacitracin is a cyclic dodecapeptide metalloantibiotic produced by some *B. licheniformis* and *B. subtilis* species (70). Bacitracin binds together with a Zn^2+^ ion to sequester the pyrophosphate moiety and first prenyl group of the UPP carrier lipid released on the outer leaflet of the membrane following the transglycosylation reaction (29). To sustain the lipid II cycle the UPP carrier must be dephosphorylated to the undecaprenyl phosphate (UP), which is required by the MraY enzyme for the synthesis of lipid I.

Consistent with its close genetic relationship to known producer species, *B. subtilis* expresses a robust intrinsic resistance to bacitracin (7, 64). The first line of defense is the BceAB ABC transporter, which dissociates the bacitracin:UPP complex in a target protection mechanism (9). The BceAB transporter is specifically induced by bacitracin through the action of the BceRS two-component system (10). The BceS sensor kinase forms a complex with the BceAB proteins to allow for a flux-sensing regulation mechanism (10, 11). The second line of defense is BcrC, a σ^M^-dependent UPP phosphatase (7). By dephosphorylating UPP to UP, BcrC converts the bacitracin target to a form no longer recognized by this antibiotic (65). However, UP is the target for amphomycin antibiotics, including the semi-synthetic derivative MX-2401 (57). Here, we identify the *ytpAB* operon as an additional contributor to intrinsic resistance to bacitracin.

YtpA is annotated as a phospholipase responsible for removal of a fatty acyl chain from phosphatidylglycerol to generate a lysophospholid species (bacilysocin) reported to have weak antibiotic activity (23). We have confirmed that *B. subtilis* does produce lysophospholipids, including LPG, detectable both in the supernatant and membrane fractions. However, YtpA is not required for LPG production (Figure 3), nor are lysophospholipid levels enhanced in a strain in which *ytpA* is induced (Supplementary Figure 3). The enzyme, presumably an A_1_ phospholipase, that is responsible for forming these species remains to be determined, and the product(s) that result from YtpA activity are also still unclear.

The *ytpAB* operon additionally encodes YtpB, an enzyme that converts heptaprenyl pyrophosphate (HPP) into the monocyclic tetraprenyl-β-curcumene in the committed step in the synthesis of the C_35_ terpenoid designated baciterpenol A (24, 25). Baciterpenol A presumably modulates membrane properties and may thereby contribute to stress resistance, but its physiological role remains poorly characterized. A *ytpB* mutant strain was previously reported to be sensitized to bacitracin, an effect was attributed to an increased accumulation of the substrate, HPP, rather than the absence of product (28). The co-regulation of YtpA and YtpB is consistent with a model in which they both function to modulate membrane properties in response to stress.

We here present evidence that *ytpA* affects membrane fluidity and interacts genetically with proteins that function in homeoviscous adaptation. Specifically, we observed a striking growth defect in Δ*bkd* Δ*ytpA* cells lacking the ability make BCFAs and desaturated phospholipids and also missing YtpA (Figure 2). This indicates that YtpA modulates membrane properties. Whether these effects are due to the modest impact that YtpA has on lysophospholipid synthesis (Figure 3), or by some other mechanism, is still unknown.

YtpA also contributes to intrinsic bacitracin resistance as revealed in cells defective in other intrinsic resistance mechanisms (Figure 6). The known mechanisms of bacitracin resistance all affect the availability of the bacitracin target UPP (BcrC) or the stability of its complex with bacitracin (BceAB) (Figure 8). In *B. subtilis*, there are two UPP phosphatases, UppP and BcrC, and at least one is required for viability (54, 55). Circumstantial evidence suggests that BcrC may be the major UPP phosphatase active on the outer leaflet of the membrane. Specifically, loss of BcrC leads to a significant increase in sensitivity to bacitracin, whereas there is no effect seen with a strain lacking *uppP*, the second UPP phosphatase. The strong additive effect on bacitracin sensitivity between *bcrC* (which increases UPP levels in the outer leaflet) and *ytpA* (Figure 6) suggests that increased membrane fluidity may facilitate UPP flipping, possibly through a spontaneous reaction or one involving an unidentified protein partner (Figure 8). In addition, YtpA may alter membrane properties that affect the function of membrane-anchored enzymes involved in cell wall synthesis.

**Figure 8.**
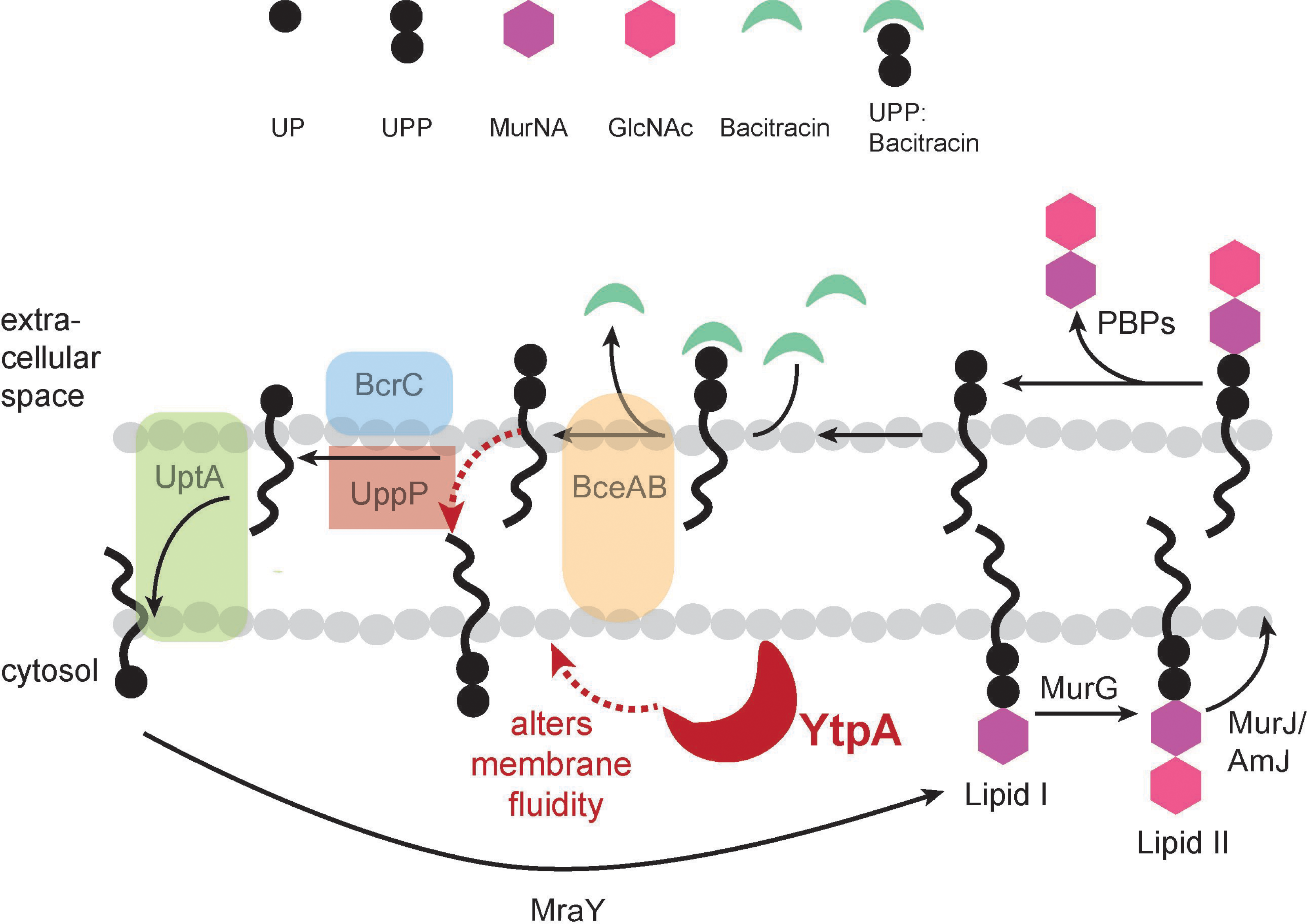
Model for lipid II recycling. Lipid-II consists of the peptidoglycan precursors bound to undecaprenyl pyrophosphate (UPP) moiety and is synthesized in the inner leaflet of the membrane. Flippases MurJ/Amj flip it to the outer leaflet where PBPs incorporate the precursors into the peptidoglycan meshwork. Subsequently, UPP is dephosphorylated to UP, and UptA (or other unidentified proteins) flips UP back to the inner membrane, where MraY initiates the incorporation of peptidoglycan precursors. Bacitracin binds to UPP in the outer membrane, inhibiting its recycling and limiting cell wall synthesis. In response to bacitracin treatment, BceAB is upregulated to remove bacitracin from UPP, and BcrC is upregulated to dephosphorylate UPP into UP, thereby eliminating the bacitracin target. In addition, YtpA, which increases membrane fluidity by an unknown mechanism, may contributes to bacitracin resistance. We speculate that YtpA may aid in flipping of UPP from the outer to the inner leaflet of the membrane.

## Materials and Methods

### Growth conditions, bacterial strains, and plasmids

All strains were cultured in lysogeny broth (LB) medium at 37 °C and aerated on an orbital shaker at 300 RPM. Before each experiment, glycerol stocks were streaked onto fresh LB agar plates and grown overnight at 37 °C. Antibiotics were used as required at the following concentrations: 100 µg/ml ampicillin; 10 µg/ml chloramphenicol; MLS (1 µg/ml erythromycin and 2.5 µg/ml lincomycin); 10 µg/ml kanamycin. Bacterial strains used in this study are listed in Table 1. Bacitracin was used as the biologically active Zn-salt (Zn bacitracin; Sigma #B5150), unless otherwise indicated. Deletion strains were created utilizing the BKK/BKE genomic library available at the Bacillus Genome Stock Center (BGSC) (71). Taking advantage of the natural competence of *B. subtilis*, all gene deletions with either a kanamycin or an erythromycin cassette were moved into the WT *B. subtilis* 168 strain. For transformation, *B. subtilis* was grown in modified competence media (MC) to stationary phase ∼0.8-1.0 OD_600nm_, incubated with desired DNA for 1-2 hours, and plated on appropriate antibiotics. Null mutations were created by removing the antibiotic resistance cassette to create clean, in-frame deletion mutants using the pDR244 plasmid, as described (71). All gene deletions were confirmed via colony PCR using designated check primers (Table S1). Genes were overexpressed ectopically at the *amyE* locus using the pPL82 plasmid (72). Ectopic overexpression from the *Pspac(hy)* promoter was induced with 1 mM IPTG. Long-flanking homology PCR was used to construct operon deletions using primers as shown in Table S1. The *Δbkd* strain contains a deletion of the entire *bkd* operon (*ptb, bcd, ipdV, bkdAA, bkdAB,* and *bkdB*), in addition to a *des* deletion (39).

**Table 1.** Strains used in this study.

| Strain Number | Genotype | Construction | Reference |
| --- | --- | --- | --- |
| 168 | *trpC2* | Lab strain | Lab stock |
| HB27141 | *trpC2 ytpA::erm* | BGSC → 168 | Lab stock |
| HB27232 | *trpC2 ytpA null* | pDR244 → HB27141 | This study |
| HB27450 | *trpC2 amyE::pSpac(hy)-ytpA* | pPL82-*ytpA* → 168 | This study |
| HB27142 | *trpC2 des::kan* | BGSC → 168 | This study |
| HB27373 | *trpC2 des::kan bkd::erm (Δbkd)* | gDNA *bkd::erm* (see primers) → HB27142 | This study |
| HB27384 | *trpC2 des::kan bkd::erm amyE::pSpac(hy)-ytpA* | pPL82-*ytpA* → HB27373 | This study |
| HB27482 | *trpC2 ytpA null des::kan bkd::erm* | gDNA *bkd::erm* (see primers) → HB27232 | This study |
| HB27246 | *trpC2 sacA::PytpAB-luxABCDE-erm* | pBS3E-lux-*PytpAB* → 168 | This study |
| HB27247 | *trpC2 sacA::PytpAB reduced-luxABCDE-erm* | pBS3E-lux-*PytpAB* reduced → 168 | This study |
| HB27260 | *trpC2 sigM::erm* | BGSC → 168 | Lab stock |
| HB27287 | *trpC2 sigM::erm sacA::PytpAB-luxABCDE-erm* | pBS3K-lux-*PytpAB* → HB27260 | This study |
| HB27272 | *trpC2 bcrC::erm* | BGSC → 168 | Lab stock |
| HB27277 | *trpC2 bcrC null* | pDR244 → HB27272 | This study |
| HB27291 | *trpC2 bcrC null sacA::PytpAB-luxABCDE-erm* | pBS3K-lux-*PytpAB* → HB27277 | This study |
| HB27271 | *trpC2 bceAB::kan* | HB0928 | Lab stock |
| HB27289 | *trpC2 bceAB::kan sacA::PytpAB-luxABCDE* | pBS3E-lux-*PytpAB* → HB27271 | This study |
| HB27249 | *trpC2 ytpB::erm* | BGSC → 168 | Lab stock |
| HB27253 | *trpC2 ytpB null* | pDR244 → HB27249 | This study |
| HB27407 | *trpC2 ytpAB::erm* | See primers | This study |
| HB27275 | *trpC2 ytpA null bcrC::erm* | gDNA HB27272 → HB27232 | This study |
| HB27360 | *trpC2 ytpA null bcrC null* | pDR244 → HB27275 | This study |
| HB27442 | *trpC2 uppP::erm* | BGSC → 168 | Lab stock |
| HB27446 | *trpC2 uppP null* | pDR244 → HB27442 | This study |
| HB27443 | *trpC2 ytpA null uppP::erm* | gDNA HB27442 → HB27232 | This study |
| HB27447 | *trpC2 ytpA null uppP null* | pDR244 → HB27443 | This study |
| HB27344 | *trpC2 uptA::erm* | BGSC → 168 | Lab stock |
| HB27393 | *trpC2 uptA null* | pDR244 → HB27344 | This study |
| HB27351 | *trpC2 ytpA null uptA::erm* | gDNA HB27344 → HB27232 | This study |
| HB27362 | *trpC2 ytpA null uptA null* | pDR244 → HB27351 | This study |
| HB27273 | *trpC2 ytpA null bceAB::kan* | gDNA HB27271 → HB27232 | This study |
| HB27490 | *trpC2 ytpA uptA bceAB::kan* | gDNA HB27271 → HB27362 | This study |

### Fluorescence Anisotropy

Fluorescence anisotropy was performed as described with modification (73). Briefly, 5 ml of cells were grown in LB medium at 37 °C with shaking to an OD_600nm_ ∼1.0 with or without 1 mM IPTG induction, where applicable. Cells were harvested and centrifuged at 2500 x g for 3 minutes. Cell pellets were washed twice and then resuspended in phosphate buffer (100 mM, pH 7.0) to OD_600nm_ 0.15. Cells were treated with 1,6-diphenyl-1,3,5-hexatriene (DPH) (Sigma) to a final concentration of 3.2 µM. An unlabeled control was also prepared. Cells were incubated in the dark in a 30 °C water bath for 30 minutes. Fluorescence anisotropy was performed with a PerkinElmer LS55 luminescence spectrometer (λ_ex_ = 358 nm, slit width = 10 nm; λ_em_ = 428 nm, slit width = 15 nm). A correction for the fluorescence intensity of unlabeled cells was performed as described (74). Data averages and standard deviations of 3 biological replicates are shown.

### Spot dilution assay

Cells were streaked onto LB agar plates and grown overnight at 37 °C. From a colony, 5 mL cells were grown in LB till ∼ 0.4 OD_600nm_. Ten-fold serial dilutions were prepared and 5 uL of the cells were plated on LB medium. Plates were allowed to air dry for 20 minutes and then incubated at 27, 37, and 42 °C. Images were captured after two days for plates incubated at 27 °C and 1 day for plates incubated at 37 and 42 °C. For the cold sensitivity assay, cells were streaked onto LB agar plates supplemented with 100 µM 2-methylbutyric acid (MB) (Sigma), and grown at 37 °C. 5 ml of cells were grown from an isolated colony in LB medium in the absence of 2-methylbutyric acid at 37 °C to ∼1.0 OD_600nm_. Cells were harvested and centrifuged at 2500 x g for 5 minutes, and the pellets were washed with an equal volume of standard lab minimal media (15 mM (NH_4_)_2_SO_4_, 0.8 mM MgSO_4_ 7H_2_O, 3.4 mM sodium citrate dihydrate, 2 mM KPO_4_, 4.2 mM potassium glutamate, 40 mM morpholinepropanesulfonic acid (MOPS), pH 7.4, 0.25 mM tryptophan, 5 µM FeSO_4_, 5 µM MnCl_2_, 2 % glucose). Ten-fold serial dilutions were performed in minimal medium, and 10 µL of cells were spotted onto minimal medium plates. Plates were allowed to air dry for 20 minutes and then incubated at 22 °C temperatures. Minimal media agar plates were either unsupplemented, supplemented with 100 µM MB, or supplemented with 1 mM IPTG. Spot dilutions were photographed every 24 hours to monitor growth. N = 3. A representative image is shown.

### LPG/LPE mass spectrometry

WT and *ytpA* deletion strains were grown in LB media till late-log phase and over-night for stationary phase cultures. For the strains harboring IPTG-inducible *ytpA*, cells were grown with or without 1 mM IPTG till late-log phase. LPG/LPE were extracted from 5 mL of cells or 1 mL of supernatant from 0.2 μm filtered media. The cells were resuspended in 0.5 mL water and 0.5 mL of cold methanol containing 1% acetic acid was added. To the 1 mL of filtered media, 1 mL of cold methanol containing 1% acetic acid was added. Samples were incubated on ice for 10 min and centrifuged at 20,000 × g for 20 min. Supernatants were dried in a speed vac concentrator and resuspended in 80% methanol containing 100 ng/mL of [d5]17-LPG.

LPG and LPE were analyzed using a Shimadzu Prominence UFLC attached to a QTrap 4500 equipped with a Turbo V ion source (Sciex). Samples were injected onto an Acquity UPLC HSS C18, 2.5mm, 3.0 x 150 mm column at 30°C (Waters) using a flow rate of 0.2 mL/min. Solvent A was 5 mM ammonium acetate + 1% formic acid, and Solvent B was 95% methanol + 5 mM ammonium acetate + 1% formic acid. The HPLC program was the following: starting solvent mixture of 35% A/65% B, 0 to 1 min isocratic with 65% B; 1 to 3 min linear gradient to 100% B; 3 to 30 min isocratic with 100% B; 30 to 32 min linear gradient to 65% B; 32 to 35 min isocratic with 65% B. The QTrap 4500 was operated in the negative mode, and the ion source parameters were: ion spray voltage, -4500 V; curtain gas, 30 psi; temperature, 500°C; collision gas, medium; ion source gas 1, 20 psi; ion source gas 2, 35 psi; declustering potential, -80 V; and collision energy, -30 V. The multiple reaction monitoring (MRM) transitions for LPG and LPE species are listed in Table S2. [d5]17-LPG was used as the internal standard. The system was controlled by the Analyst software (Sciex) and analyzed with MultiQuant 3.0.2 software (Sciex). Peaks corresponding to individual LPG species were quantified relative to the internal standard.

### Phosphatidylglycerol (PG) mass spectrometry

WT and *ytpA* deletion strains were grown in LB media till late-log phase. Lipids were extracted from 5 mL of culture by the Bligh and Dyer method. Lipid extracts were resuspended in chloroform/methanol (1:1). PG was analyzed using a Shimadzu Prominence UFLC system attached to a QTrap 4500 equipped with a Turbo V ion source (Sciex). Samples were injected onto an Acquity UPLC BEH HILIC, 1.7 µm, 2.1 × 150 mm column (Waters) at 45 °C with a flow rate of 0.2 mL/min. Solvent A was acetonitrile, and solvent B was 15 mM ammonium formate, pH 3. The HPLC program was the following: starting solvent mixture of 96% A/4% B; 0 to 2 min, isocratic with 4% B; 2 to 20 min, linear gradient to 80% B; 20 to 23 min, isocratic with 80% B; 23 to 25 min, linear gradient to 4% B; 25 to 30 min, and isocratic with 4% B. The QTrap 4500 was operated in the Q1 negative mode. The ion source parameters for Q1 were as follows: ion spray voltage, −4500 V; curtain gas, 25 psi; temperature, 350 °C; ion source gas 1, 40 psi; ion source gas 2, 60 psi; and declustering potential, −40 V. The system was controlled and analyzed by the Analyst software (Sciex).

The samples were introduced to the QTrap 4500 by direct injection to perform product scans to verify the fatty acids present in a particular PG molecular species along with the positional distribution of the fatty acids. The ion source parameters for negative mode product scan were as follows: ion spray voltage, −4500 V; curtain gas, 10 psi; collision gas, medium; temperature, 270 °C; ion source gas 1, 10 psi; ion source gas 2, 15 psi; declustering potential, −40 V; and collision energy, −50 V.

### Luciferase reporter construction and measurement

Luciferase reporters were constructed by inserting designated promoters into the multiple cloning site of pBS3E*lux*, or pBS3K*lux* (75) and transformed into *B. subtilis* using natural competence as described above. For luciferase measurements, strains were grown in LB medium at 37 °C to ∼0.4 OD_600nm_. 2 μl of culture was inoculated into 99 µl of fresh LB medium in a 96 well plate. Where applicable, cultures were treated with 0.005 µg/ml cefuroxime. The concentration of bacitracin used varied depending on the strains and has been mentioned in the figure legend. The plate was incubated at 37 °C with orbital shaking in a Synergy H1 plate reader (BioTek Instruments, Inc,) and OD_600nm_ and luminescence was measured every 6 minutes. Relative light units (RLU) for promoter activity were determined by luciferase intensity normalized for cell density (OD_600nm_). Data shown is the representative average and standard deviations of 3 biological replicates.

### Growth kinetics assay

From a single colony, cells were grown in 5 ml LB medium at 37 °C with shaking to OD_600nm_ ∼ 0.4-0.5. 1 μL of culture was added to 199 μL of fresh LB medium in a 100-well Honeycomb plates (Steri). Where applicable, cells were treated with sub-lethal concentrations of bacitracin as determined by the relative bacitracin sensitivity of each strain. The OD_600nm_ of each well was measured at 37°C with shaking in a Bioscreen C Pro growth analyzer (Growth Curves USA, NJ) every 30 minutes for 24 hours. Data shown are representative plots and standard deviations are from three biological replicates.

### Real-time PCR

Gene expression was determined by real-time PCR using primers mentioned in Table S1. Cultures were grown up to an OD_600nm_ of ∼0.4. RNA was purified from 1.5 mL of cells using the RNeasy kit from Qiagen as per the manufacturer’s instructions. The isolated RNA was then given a DNase treatment with a Turbo DNA-free kit (Invitrogen, AM1907). Approximately 15 μg of RNA was incubated with 2 μL of DNase and 2 μL of buffer at 37°C for 15 min, followed by a 5-min incubation with the DNase-inactivating agent. The samples were then centrifuged at 8,000 rpm for 3 min, and the supernatant was collected in a fresh microcentrifuge tube. cDNA was prepared with 2 μg of the treated RNA in 20 μL total volume of reaction mix using a high-capacity cDNA reverse transcription kit from Applied Biosystems (4368814). The cDNA was further diluted 1:10 to obtain a final concentration of 10 ng/μL. Gene expression levels were measured using 10 ng of cDNA, 0.5 μM gene specific primers, and 1× SYBR green master mix (Applied Biosystems, A25742). The *gyrA* gene was used as an internal control.

## Acknowledgements

This work was supported by National Institutes of Health grants R35 GM122461 (J.D.H.), GM034496 (C.O.R.), Cancer Center Support Grant CA21765, and ALSAC, St. Jude Children’s Research Hospital. The authors thank Dr. David Rudner for providing MX-2401 and Karen Miller for the preparation of samples for the lipidomic experiments. The content is solely the responsibility of the authors and does not necessarily represent the official views of the National Institutes of Health.

